# BMP signaling: A significant player and therapeutic target for osteoarthritis

**DOI:** 10.1101/2021.03.01.433366

**Authors:** Akrit Pran Jaswal, Bhupendra Kumar, Anke J. Roelofs, Sayeda Fauzia Iqbal, Amaresh Kumar Singh, Anna H.K. Riemen, Hui Wang, Sadaf Ashraf, Sanap Vaibhav Nanasaheb, Nitin Agnihotri, Cosimo De Bari, Amitabha Bandyopadhyay

**Affiliations:** Department of Biological Sciences and Bioengineering, Indian Institute of Technology Kanpur, Kanpur-208016, Uttar Pradesh, India; The Mehta Family Centre for Engineering in Medicine, Indian Institute of Technology Kanpur, Kanpur, Uttar Pradesh, India; Arthritis and Regenerative Medicine Laboratory, Aberdeen Centre for Arthritis and Musculoskeletal Health, Institute of Medical Sciences, University of Aberdeen, Aberdeen AB25 2ZD, UK; Department of Zoology, Banaras Hindu University, Varanasi-221005, Uttar Pradesh, India

**Keywords:** BMP, Osteoarthritis, articular cartilage, local inhibition, LDN-193189

## Abstract

**Objective:** To investigate the role of BMP signaling in osteoarthritis' pathogenesis and propose a disease-modifying therapy for OA.

**Methods:** C57BL6/J mouse line was used to perform ACLT surgery at P120 to study the expression pattern of the BMP signaling readout pSMAD1/5/9. To investigate whether activation of BMP signaling is sufficient and necessary to induce osteoarthritis, we have used conditional GOF and LOF mouse lines in which BMP signaling can be activated or depleted, respectively, upon intra-peritoneal injection of tamoxifen. Finally, we locally inhibited BMP signaling through intra-articular injection of LDN-193189 pre- and post-onset surgically induced OA. Most of the analysis has been done through immunohistochemistry, histopathological staining, and micro-CT to evaluate the status of the pathogenesis of the disease.

**Results:** We observed concomitant activation of BMP signaling, as judged by pSMAD1/5/9 immunoreactivity in the articular cartilage, upon induction of osteoarthritis with simultaneous depletion of SMURF1, an intra-cellular BMP signaling inhibitor in articular cartilage. Even without surgical induction of osteoarthritis, only BMP gain-of-function mutation induces OA in mouse articular cartilage. Also, genetic, or pharmacological inhibition of BMP signaling offered significant protection against OA pathogenesis. Interestingly, post-onset of the disease, inhibition of BMP signaling by intra-articular injection of LDN-193189 retarded OA progression with a significant reduction in inflammatory markers.

**Conclusion:** Our study demonstrated that BMP signaling plays an essential role in the pathogenesis of OA and that local BMP inhibition can be an effective therapeutic strategy to mitigate osteoarthritis.

## Introduction

Osteoarthritis (OA) is a painful, debilitating musculoskeletal disorder with a profound socioeconomic burden and is the primary cause of locomotive disability affecting millions of people worldwide(*1*–*3*). The alarmingly increasing prevalence of OA is exacerbated further as no therapy exists to manage OA except for symptomatic treatment with anti-inflammatory drugs or surgical intervention in late stage disease(*4*). It is imperative, therefore, to discern the molecular basis of pathogenesis of OA to develop a disease modifying therapy. Articular cartilage, the tissue affected in OA, is a lubricated, avascular, alymphatic and aneural that lines the ends of the bones at the joints. During OA, the joint surface undergoes a slew of changes characterised by loss of cartilage proteoglycans, hypertrophy of chondrocytes, angiogenesis, osteophyte formation, and ultimately failure of joint function(*4*). The cellular and molecular changes of the joint cartilage during the onset and progression of OA closely resemble the steps of endochondral ossification, the developmental process by which long bones form within cartilage anlagen(*5*, *6*).

During endochondral ossification, most of the initial cartilage mass in an appendicular skeletal element is replaced by newly formed bone, except for the cartilage at the termini. The cartilage that is replaced by bone is referred to as the transient cartilage, while the cartilage at the terminal ends is referred to as the joint or articular or permanent cartilage(*7*). During transient cartilage differentiation, type II collagen (Col2a1-expressing cartilage cells undergo a series of changes. These cells undergo pre-hypertrophic differentiation wherein they express Indian hedgehog (IHH), subsequently the transition from pre-hypertrophy to hypertrophy is marked by the expression of type X collagen (ColX). The hypertrophic cells are infiltrated by blood vessels. This is followed by matrix remodelling, where enzymes viz. MMP-13 and ADAMTS-5, degrade the existing collagen matrix and a new matrix, rich in type I collagen (ColI), is synthesised and bone formation is accomplished(*8*, *9*).

Ray et *al*. discovered a zone of Col2a1-expressing bipotential proliferating cells known as the Distal Proliferative Zone (DPZ) within a developing appendicular skeletal element. The DPZ cells under the influence of BMP signaling undergo transient cartilage differentiation, whereas when exposed to Wnt signaling they undergo joint cartilage differentiation(*1*). Some of the molecules involved in transient cartilage differentiation, viz. MMP-13, ADAMTS-5, and VEGF-A, are reported to be associated and/or necessary for the pathogenesis of OA (*10*–*16*).

Previous literature suggests that ectopic activation of BMP signaling in developing cartilage or presumptive joint sites, either by overexpression of BMP ligands(*1*, *17*) or misexpression of constitutively active BMP receptors(*18*), results in transient cartilage differentiation at the expense of joint cartilage. A surge in BMP2 and BMP4 ligands was reported in human articular cartilage having a moderate to severe form of osteoarthritis(*19*). Blocking BMP signaling inhibits chondrocyte hypertrophy and mineralization indicates it is important player in terminal differentiation of BMSCs (*20*).

Additionally, Noggin administration in an ACLT induced OA model inhibits OA progression by inhibiting IL-1β and BMP-2(*21*). A recently published in-vitro study indicates reduction of chondrocyte hypertrophy after BMP receptors were inhibited using LDN-193189(*22*).Immobilisation of developing embryonic limbs leads to ectopic differentiation of transient cartilage at the cost of articular cartilage. Moreover, it was shown that immobilization induced OA leads to ectopic upregulation of BMP signaling within the sub-articular cartilage domain where cartilage precursors are normally exposed only to Wnt signaling (*23*). Recently, it was also demonstrated that pharmacological inhibition of BMP signaling promotes articular cartilage differentiation in hMSC derived chondrocytes and allows the cells to maintain an articular chondrocyte phenotype for longer a duration of time upon implantation in mice(*2*), suggesting that an embryonic paradigm of spatial restriction of BMP signaling is needed for differentiation and maintenance of the articular cartilage phenotype. However, few studies indicate BMPs have an anabolic effect on articular cartilage integrity(*24*).

Taken together, we hypothesised that BMP signaling-induced transient cartilage differentiation within the adult articular cartilage domain is the molecular basis of the pathogenesis of OA. In this study, we tested this hypothesis with conditional gain-and loss-of-function mouse mutants of BMP signaling in conjunction with a surgically induced model of OA. Our findings in the mouse model are further supported by data obtained from osteoarthritic human cartilage specimens, wherein we found evidence of active BMP signaling in the joint cartilage. Moreover, our data indicates that pharmacological inhibition of BMP signaling in the synovial joint may serve as an effective disease modifying therapy for OA.

## Materials and Methods

Details of methodology is given in supplementary section.

### Generation of mice lines

All animals were housed, bred, and maintained in Central Experimental Animal Facility (CEAF) of Indian Institute of Technology Kanpur, India. All experiments were performed in accordance with the guidelines of the Institutional Animal Ethics Committee (IAEC) as well as under the aegis of the Centre for Purpose of Control and Supervision of Experiments on Animals (CPCSEA), Government of India under protocols IITK/IAEC/2013/1002; IITK/IAEC/2013/1015; IITK/IAEC/2013/1040 and IITK/IAEC/2022/1166. We obtained B6By/J wild type mice, *TgTgCol2a1-Cre-ERT2* (46) and *ROSA26 mT/mG* (47) strains from Jackson Laboratories, USA; *pMes-caBmpr1a* mice as gift from Prof. YiPing Chen at Tulane University, USA; Bmp2c/c; Bmp4c/c mice from Prof. Clifford Tabin at Harvard Medical School, USA. For BMP signaling gain-of-function, pMes-caBmpr1a mice were crossed with TgTgCol2a1-Cre-ERT2 mice to generate *pMes-caBmpr1a; TgCol2a1-Cre-ERT2. Bmp2c/c; Bmp4c/c* (29) animals were crossed with *TgTgCol2a1-Cre-ERT2* to generate *Bmp2c/c; Bmp4c/c; TgTgCol2a1-Cre-ERT2* for BMP loss-of-function mutation.

### Anterior Cruciate Ligament Transection (ACLT)

ACLT surgeries were conducted at P120 left limb of B6By/*J* wild type male mice and harvested at post ACLT day 07, 14, 21, 28, and 56 for molecular and histological analysis. ACLT in Bmp2c/c; Bmp4c/c; *TgTgCol2a1-Cre-ERT2* mice line for BMP2/4 loss-of-function conducted at P84 after 14^th^ day of TAM injection. Animals were anesthetized using isoflurane and fallowed standard protocol during surgical procedure (28).

### Human sample collection

Osteoarthritic cartilage from 6 patients and non-osteoarthritic cartilage from one patient undergoing knee joint excision for malignancy at a site not involving sampled area, were obtained after informed consent and in accordance with the relevant guidelines and regulations, with approval from the NHS Grampian Biorepository Tissue Bank Committee, UK.

### OARSI scoring

The OARSI scores were calculated using the recommended guidelines for assessment of osteoarthritic severity in small animals (mice)(*26*, *27*).

### Micro-Computed Tomography (μCT)

Images were reconstructed and analysed using NRecon v1.6 and CTAn 1.16.8.0, respectively. Fixed tissues were taken in 5ml microfuge tube in hydrated condition and imaged using high resolution μCT (Skyscan 1172).

### Statistics

Graph Pad Prism 8.0.2 software was used to ascertain statistical significance of the OARSI histopathological scoring data. One-way ANOVA with post-hoc analysis (Dunnett’s test) and Unpaired t- test was performed to calculate the statistical significance and interdependence between different experimental groups. The results were plotted using Graph Pad Prism 8.0.2 software. The error bars represent Standard Deviation (S.D.)

## RESULTS

### 1. Overexpression of BMP signaling in adult joint cartilage is sufficient to induce the development of an OA-like phenotype in mice

To examine whether overexpression of BMP signaling in the articular cartilage is sufficient to induce osteoarthritis like changes in adult mice, we activated BMP signaling in postnatal cartilage at P70 by injecting tamoxifen intraperitoneal cavity of *pMes-caBmpr1a; TgCol2a1-Cre-ERT2* mouse (Fig.1A) (*Referred to as induction from here on*). Seven days of over-expression of *caBmpr1a* in adult mouse articular cartilage, ectopic activation of canonical BMP signaling, as assessed by immunoreactivity towards phosphorylated SMAD1/5/9, was observed, and it peaks after two weeks (Fig.1C′-C⁗). Expression of IHH, which marks a pre-hypertrophic state of cartilage, was observed within 7 days of induction and by 14^th^ day after induction, IHH expression has been reduced (Fig.1D-D″). ColII expression pattern depletes on the 14th post-induction day and reaches a nadir on the 56th post-induction day (Fig.1E-E⁗). The ColX expression, indicative of cartilage hypertrophy, was observed 14 days after induction, with the largest extent of hypertrophy occurring 56 days later (Fig.1F-F⁗). Embryonic (*23*, *28*), as well as adult articular cartilage cells (*2*), are proliferation deficient while transient cartilage cells are proliferative(*1*). In our experiments, we observed that along with other markers of transient cartilage differentiation, proliferation was also stimulated in the adult mouse articular cartilage after activation of BMP signaling. BrdU uptake increased in joint cartilage 7 days after induction reaching a peak on 14^th^ day of induction (Fig.1G-G″). Safranin O/Fast Green staining revealed a loss of proteoglycan staining in multiple zones with vertical clefts in the articular cartilage (Fig. 1H-1H′). OARSI scoring for integrity of articular cartilage indicates the severity of loss of articular cartilage in TAM injected versus control samples (Fig.1I). A similar trend to transient cartilage differentiating is indicated by quantification of ColII and ColX expression in control tissues vs samples injected with TAM (Fig.1J and Fig.1K).

**Fig. 1.**
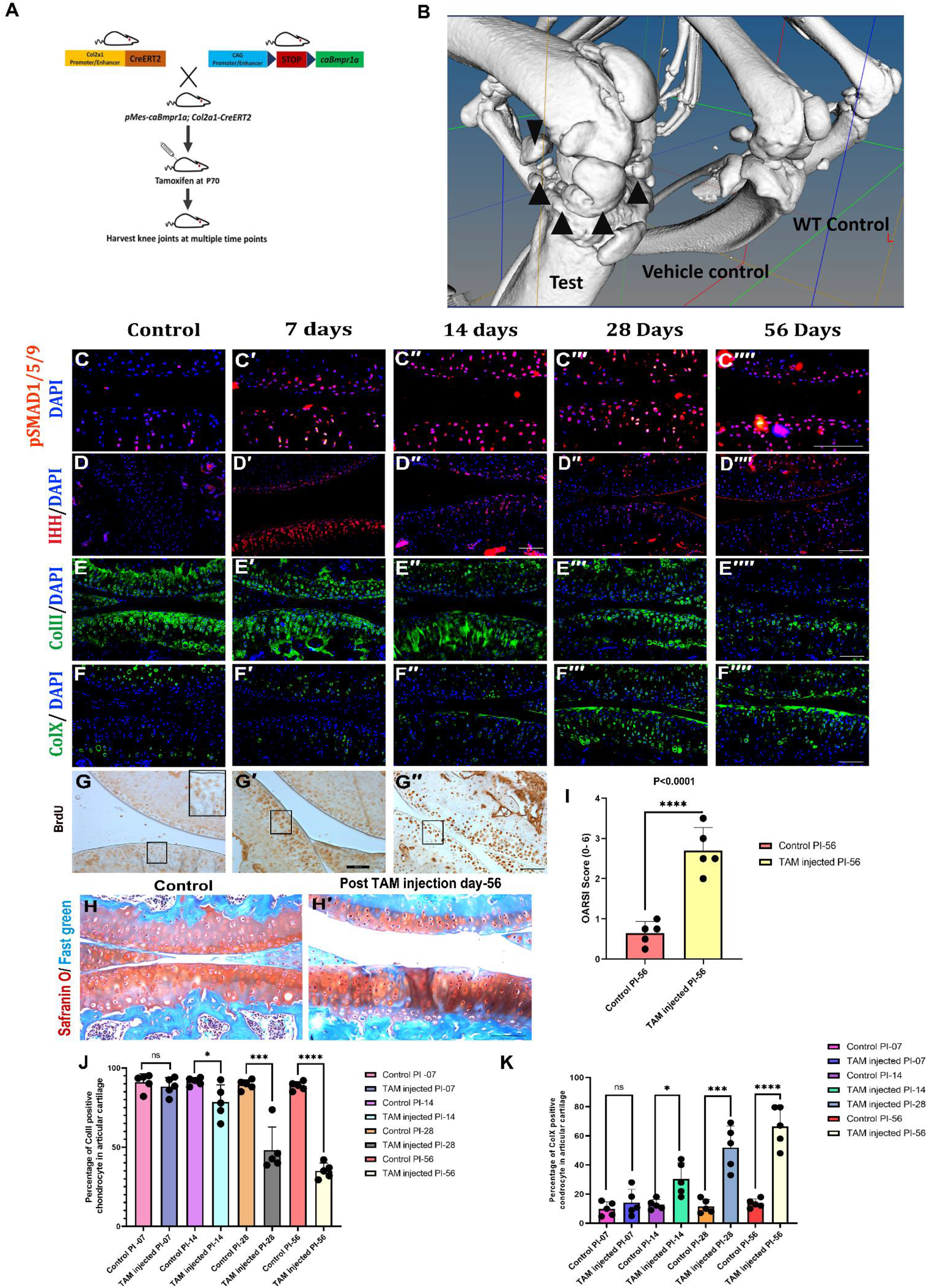
Overexpression of BMP signaling in adult joint cartilage is sufficient to induce OA development. **(A)** Schematic for generation of *pMes-caBmpr1a; TgCol2a1-Cre-ERT2* mice and mis-expression of constitutively active *Bmpr1a* in the adult cartilage by injecting tamoxifen (TAM) intraperitoneally at P70. **(B)** 3-D rendering of μCT scan at 40μm resolution in wildtype (WT) control, vehicle control and TAM injected knee joint at 180 days post induction, black arrows show osteophytes **(C-F⁗)** Longitudinal sections through the adult knee joints of vehicle control (C-H) and mice 7 days (C′-G′), 14 days (C″-G″), 28 days (C‴, F‴), 56 days (C⁗-F⁗) post induction by TAM injection. Immunoreactivity for pSMAD1/5/9 (C-C⁗), IHH (D-D⁗), ColII (E-E⁗) and ColX (F-F⁗). (**G-G″)** BrdU incorporation 7 days (G′) and 14 days (G″) after TAM injection. (**G**) Vehicle control. (**H-H′)** Safranin O staining in vehicle control (H) and TAM injected knee joints at 180 days (H′) post induction. (**I)** Statistical analysis by Unpaired t-test of OARSI scores at post TAM injection day 56 with control, p<0.0001 (****).**(J)** Quantification data for ColII, Unpaired t- test was performed to compare the means of stage matched control vs post injected (PI) TAM test animals at different time points, Control vs Test-PI day 7, p=0.4573 (ns), Control vs Test, PI day 14, p=0.0301(*), Control vs Test, PI day 28, p=0.0003 (***), Control vs Test, PI day 56, p<0.0001(****). **(K)** Quantification data for ColX, Unpaired t- test was performed to compare the means of stage matched control vs post injected (PI) TAM test animals at different time points, Control vs Test-PI day 7, p=0.3731 (ns), Control vs Test, PI day 14, p=0.0101 (*), Control vs Test, PI day 28, p=0.0004 (***), Control vs Test, PI day 56, p<0.0001 (****). n=5 per group. Scale bar = 100μm

Besides the molecular signatures, Micro CT imaging of hind limbs revealed extensive osteophyte (indicated by black arrow) formation upon ectopic activation of *Bmpr1a* in the articular cartilage (Fig.1B). Taken together, these observations indicate that ectopic activation of BMP signaling is sufficient to induce the development of an OA like phenotype inn adult mice.

### 2. BMP signaling induced transient cartilage differentiation is necessary for the pathogenesis of OA

Next, we investigated the necessity of BMP signaling in the development of the osteoarthritic phenotype. It has been previously reported that levels of BMP-2 ligands are elevated in synovial fluid from OA patients and BMP receptor localisation is associated with OA severity (*19*, *29*). We performed anterior cruciate ligament transection (ACLT) to induce OA in mice and examined BMP signaling readout pSMAD1/5/9 in knee articular cartilage every week following ACLT. (*30*, *31*)... In comparison to sham operated knees (Fig. S1A) or 7 days post ACLT (Fig. S1B), we found increased pSMAD1/5/9 immunoreactivity 14 days after ACLT (Fig. S1B′), which lasted until 56 days after ACLT (Fig. S1B″, Fig. S1B‴, and Fig. 2B′). we also found increase in In line with our findings in the context of ectopic BMP signaling activation, we found increased of BrdU uptake in the articular cartilage of mice fallowing ACLT (Fig. S1C and S1D-D‴). In order to prevent activation of BMP signaling post ACLT, we used a previously described *Bmp2/4* double conditional knockout strain(*32*). We used *Bmp2*^*c/c*^; *Bmp4*^*c/c*^; *TgCol2a1-Cre-ERT2* mice line to inject tamoxifen intraperitoneally at P70 thereafter ACLT was performed at P84 (Fig. S2A and Fig. 2A).

**Fig. 2.**
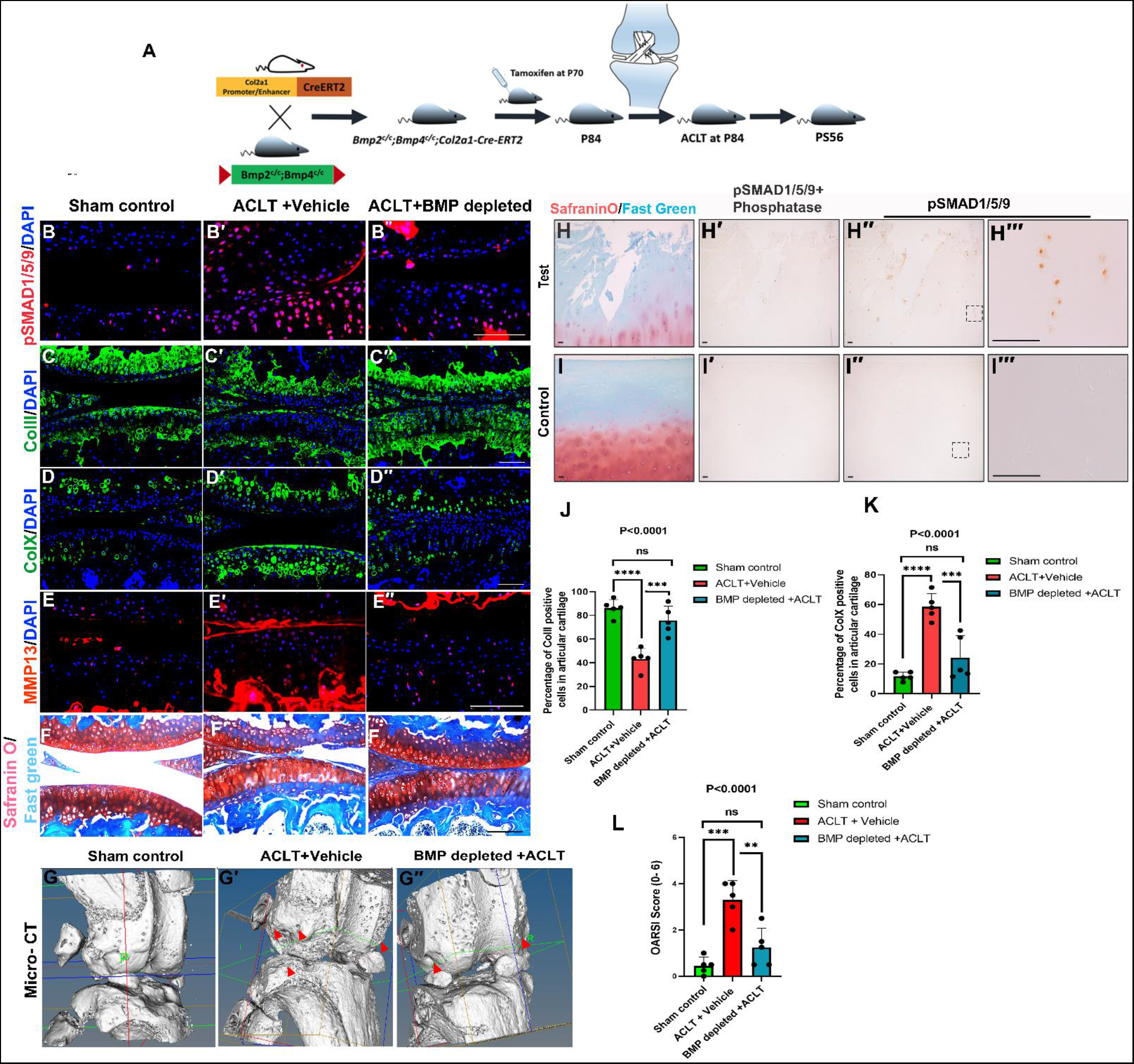
BMP signaling induced transient cartilage differentiation is necessary for the pathogenesis of OA. **(A)** Schematic representation depicting the generation of *Bmp2*^*c/c*^; *Bmp4*^*c/c*^; *TgCol2a1-Cre-ERT2* and the regimen for depletion of BMP signaling by administration of tamoxifen followed by ACLT. **(B-F″)** Longitudinal sections through the knee joints of sham (B-F), “ACLT + vehicle” control (B′-F′) and “BMP depletion + ACLT” (B″-F″) mice at 56 days post-surgery (PS56). Immunoreactivity for pSMAD1/5/9 (B-B″), ColII (C-C″), ColX (D-D″), MMP-13 (E-E″). (F-F″) Safranin O staining. (**G-G″**) 3-D rendering of μCT at PS56. Red arrowheads indicate osteophytes, surface roughness, and damage. n=5 per time point per group. Scale bar = 100μm. **(H-I‴)** Histological sections of knee articular cartilage from OA patients (n=6) (J-J‴), and a patient without known history of knee OA (n=1) (I-I‴). (H, I) Safranin O/Fast Green staining of OA (H) and normal (I) cartilage. Immunoreactivity for pSMAD1/5/9 with (H′, I′) or without phosphatase pre-treatment to verify antibody specificity (H″, H‴, I″, I‴), of OA (H′-H‴) and normal (I′-I‴) cartilage. (H‴, I‴) Higher magnification view of the marked regions in H″ and I″. **(J)** Quantification data for ColII, one way ANOVA was performed along the three sets and p<0.0001(****). We compared the means of sham control vs ACLT+vehicle; p<0.0001 (****), sham control vs BMP depleted+ACLT; p=0.2319 (ns) and ACLT+vehicle vs. BMP depleted+ ACLT p=0.0005(***) **(K)** Quantification data of ColX., one way ANOVA was performed along the three sets and p<0.0001(****) the means of sham control vs ACLT+vehicle; p<0.0001 (****), sham control vs BMP depleted+ACLT; p=0.1595 (ns) and ACLT+vehicle vs. BMP depleted+ ACLT p=0.0004(***) **(L)** OARSI score, one-way ANOVA was performed, p=0.0001(***), the means of sham control vs ACLT+vehicle; p=0.0001 (***), sham control vs BMP depleted+ACLT; p=0.2195 (ns) and ACLT+vehicle vs. BMP depleted+ ACLT p=0.0018(**) Scale bar = 100μm. The panels where *Bmp2/4* depleted animals were subjected to ACLT are marked as “BMP depletion + ACLT”. Vehicle injected animals were used as genotype controls (“ACLT + Vehicle”. “Sham” refers to *Bmp2*^*c/c*^; *Bmp4*^*c/c*^; *TgCol2a1-Cre-ERT2* animals which underwent sham surgery without ACLT.

As expected, after ACLT, pSMAD1/5/9 immunoreactivity was minimal in articular cartilage of Bmp2/4-depleted animals. (Fig. S2B″ and Fig. 2B″). Distribution and abundance of ColII was significantly preserved in *Bmp2*/*4* depleted animals even after 56 days of ACLT (Fig. S2C-C″ and Fig. 2C-C″). Chondrocyte hypertrophy, as assessed by ColX immunoreactivity (Fig. 2D-D″) as well as expression of MMP-13 (Fig. 2E-E″), a key matrix remodelling enzyme, were remarkably elevated after 56 days of ACLT (Fig. 2D′ and Fig. 2E′). However, the depletion of Bmp2/4 shielded from upregulation and allowed the maintenance of a ColX. (Fig. 2D″) and MMP-13 (Fig. 2E″) which were almost comparable to that of sham (Fig. 2D and Fig. 2E) Articular cartilage loss was observed in ACLT specimens as measured by Safranin O/Fast green staining, these changes were minimal in BMP ligand depletion specimen (Fig. 2F-F″). Micro-computed tomography (μCT) structural examination revealed that the "ACLT + Vehicle" group had extensive damage to articular surfaces (roughness) as well as osteophyte formation (marked by red arrows) (Fig.2G′). However, these changes were minimal in BMP ligands were depleted specimen, the severity and extent of these changes were minimal, and were significantly (Fig. 2G″), and comparable to sham operated group (Fig.2G), indicating that cartilage protection was provided. Quantification of ColII and ColX in the ACLT+BMP depleted group reveals significant similarity with the Sham control (Fig. 2J & 2K). OARSI scoring indicates significant protection of articular cartilage integrity in the BMP depleted +ACLT group compared to the ACLT+vehicle group (Fig. 2L)

To ascertain the clinical relevance of these findings, we examined both osteoarthritic and non-osteoarthritic human articular cartilage. pSMAD1/5/9 immunoreactivity was found in all zones of osteoarthritic cartilage from patients who had arthroplasty (Fig. 2H″, 2H‴), whereas human cartilage from a donor with no known history of OA showed no detectable pSMAD1/5/9 immunoreactivity (Fig. 2I″, 2I‴). There was no pSMAD1/5/9 immunoreactivity in phosphatase-treated osteoarthritic cartilage (Fig. 2H′, 2I′)

### 3. Local pharmacological inhibition of BMP signaling halts the progression of osteoarthritic changes

In order to determine if local inhibition of BMP signalling after ACLT would slow the progression of osteoarthritis in mice, LDN-193189, a well-known dorsomorphin derivative and BMP signalling inhibitor, was administered in the joint cavity. (*33*–*35*). LDN-193189 activity was assayed using the BRITER (BMP Responsive Immortalized Reporter) cell line. (*25*) (see Materials and Methods). LDN-193189 inhibited BMP signaling in the BRITER cell line at concentrations as low as 100 nM (Fig. S3).

Considering possible dilution and volume loss of LDN-193189 during the injection, we used 6μl of 10 μM (in 3% w/v 2-hydroxypropyl-β-cyclodextrin in PBS) of LDN-193189 for intra-articular injection to inhibit BMP signaling fallowing ACLT. Seven consecutive doses of LDN-193189 was given starting from 14^th^ to 21^st^ day post-surgery and tissue were harvested at 28 days post-surgery (Fig. 3A).

**Fig. 3.**
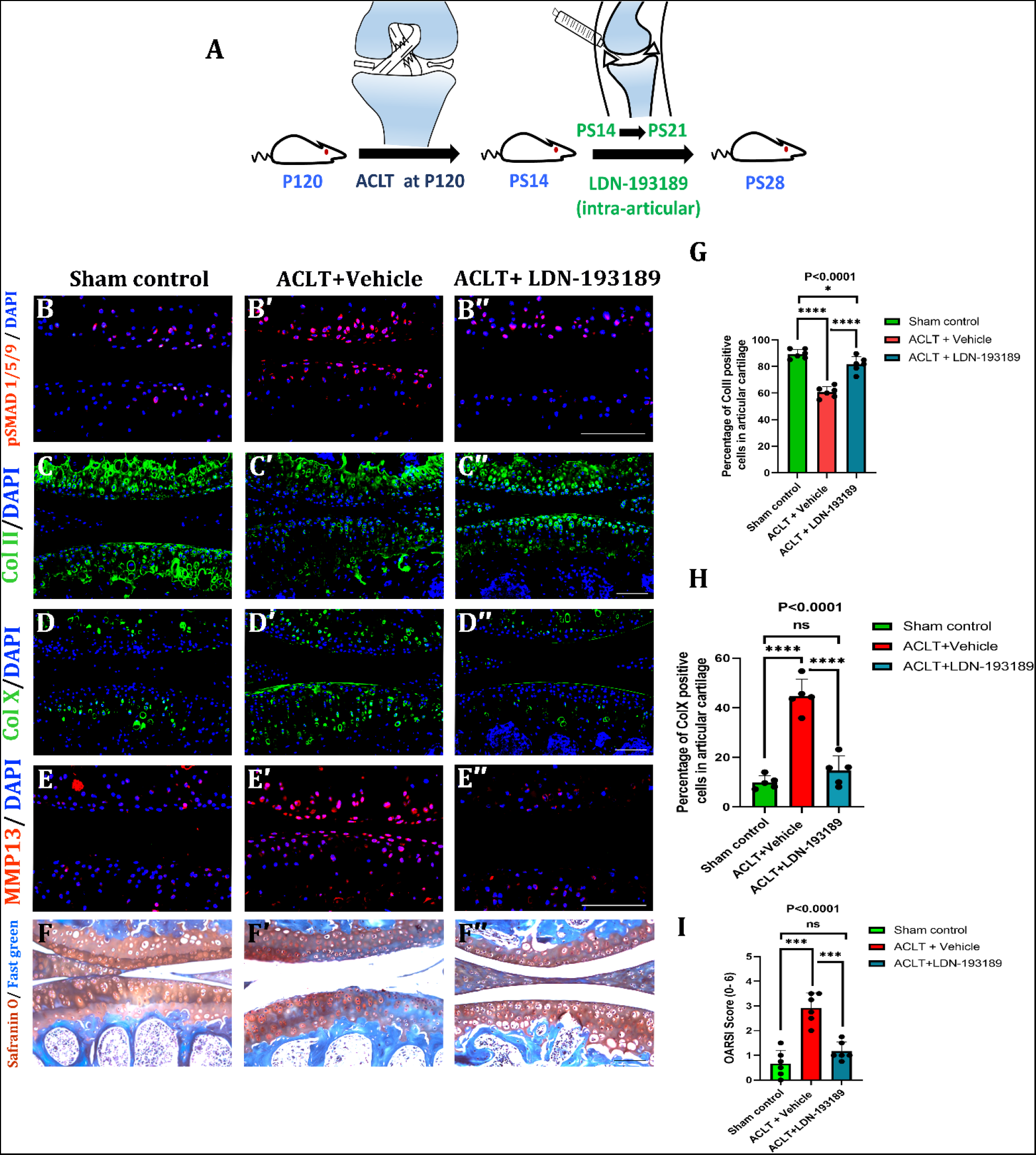
Local pharmacological inhibition of BMP signaling halts the progression of osteoarthritic changes. **(A)** Schematic for local inhibition of BMP signaling using LDN-193189 in surgically induced OA in wildtype mice. **(B-F″)** Longitudinal sections through the knee joints of sham (B-F), “ACLT + vehicle” control (B′-F′) and “ACLT + LDN-193189” (B″-F″) mice at 28 days post-surgery (PS28). Immunoreactivity for pSMAD1/5/9 (B-B″), ColII (C-C″), ColX (D-D″), MMP-13 (E-E″). (F-F″) Safranin O staining. **(G)** Quantification data for ColII, one way ANOVA was performed along the three sets and p<0.0001(****). The comparison of Sham control vs ACLT+vehicle; p<0.0001 (****), Sham control vs ACLT+ LDN-193189; p=0.0263 (*) and ACLT+vehicle vs. ACLT+ LDN-193189 p<0.0001(****). **(H)** Quantification data for ColX, one way ANOVA was performed along the three sets and p<0.0001(****). The comparison of Sham control vs ACLT+vehicle; p<0.0001 (****), Sham control vs ACLT+ LDN-193189; p=0.3897 (ns) and ACLT+vehicle vs. ACLT+ LDN-193189 p<0.0001(****). **(I)** OARSI score, one-way ANOVA was performed, p<0.0001(****), the comparison of means of Sham control vs ACLT+vehicle; p=0.0001 (****), Sham control vs ACLT+LDN-193189; p=0.2460 (ns) and ACLT+vehicle vs. ACLT+LDN-193189 p<0.0001(****); Scale bar = 100μm, n=5 per group.

We found local inhibition of BMP signaling significantly abrogates OA like changes fallowing ACL transection in mice. The pSMAD1/5/9 positive cells were found in zones of articular cartilage in vehicle administered ACLT knee joints (Fig.3B′) while lesser immunoreactivity to pSMAD1/5/9 was observed in articular cartilage of LDN-193189 treated set post ACLT (Fig. 3B″) and the sham operated knee cartilage (Fig.3B), The immunoreactivity against ColII in LDN-193189 treated knee joints was like sham operated (Fig. 3C and 3C″) as compared to ACLT+vehicle group (Fig.3C′) suggesting reduced depletion of ColII post-surgery and protection of cartilage. The hypertrophy of cartilage cells was found to be limited to the calcified zones, with minimal ColX immunostaining in the articular cartilage of LDN-193189 treated ACLT induced OA mice (Fig. 3D″), similar to the sham set (Fig. 3D), whereas vehicle injected animals showed extensive hypertrophy throughout the cartilage matrix (Fig. 3D′). Similarly, MMP-13 levels in articular cartilage were found to be significantly reduced after intra-articular administration of LDN-193189 (Fig.3E″), whereas a global upregulation of MMP-13 was observed in vehicle-injected knee joints (Fig.3E′). Proteoglycan depletion and cartilage damage were found to be minimal in the tibial surface of LDN-193189 injected patients (Fig. 3F″) when compared to the vehicle injected group (Fig. 3F′), and cartilage integrity was found to be comparable to sham operated knees (Fig. 3F). ACLT+LDN-193189 injected samples had similar ColII quantification data to sham operated controls. However, it was significantly lower in ACLT+vehicle injected samples (Fig.3G). Similarly, quantitative data for ColX expression in ACLT+LDN-193189 injected samples was similar to sham operated samples and significantly lower than ACLT+vehicle injected samples (Fig.3H). Moreover, OARSI scoring of cartilage revealed a significantly attenuated osteoarthritic-like phenotype in the LDN-193189 treated group as compared to the vehicle-treated ACLT group, and it was similar to the sham-operated group (Fig. 3I).Taken together, these findings suggest that *in situ* inhibition of BMP signaling in articular cartilage is sufficient to prevent the phenotypic and molecular changes associated with the development and progression of OA in a surgically induced osteoarthritic mouse model.

### 4. Inhibition of BMP signaling post-onset of OA attenuates disease severity

*In situ* inhibition of BMP signaling before the onset of OA following ACL transection in mice retards the progression of OA. However, in a clinical setting, patients report to the clinic after the disease has set in. We therefore investigated if local inhibition of BMP signaling can mitigate the severity of osteoarthritic changes even after the disease has set in. For this purpose, seven consecutive intra-articular LDN-193189 injections were administered starting on post-surgery day 35 and finishing on post-surgery day 42. The knees were harvested at post-surgery day 56 (Fig.4A). In contrast to the vehicle-treated knee joints (Fig. 4B′), ColII positive cells were found throughout the articular cartilage in the LDN-193189-treated samples (Fig. 4B″), which is very similar to the sham-operated group (Fig. 4B). The vehicle-treated group had significantly higher ColX and MM13 immunoreactivity than the LDN-193189-injected and sham-operated groups (Fig.4C-4C″ and Fig.4D-4D″ respectively). Articular cartilage integrity, as determined by Safranin O staining, was preserved in LDN-193189 treated knee joints and was comparable to sham operated knees (Fig.4E and 4E″), whereas vertical cleft and articular cartilage loss were observed in vehicle treated ACLT knee joints (Fig. 4E′). The μCT imaging reveals that cartilage surface erosion was reduced in the LDN-193189-treated knees compared to the vehicle-injected knees. (Fig. 4F-4F″; red arrow marks osteophytes). The quantification of ColII expression was significantly higher in the case of ACLT+LDN-193189 injected samples than vehicle injected control and it was close to sham operated samples (Fig. 4G). Similarly, quantified data for ColX immunoreactivity was higher in vehicle injected samples while it was significantly reduced in LDN-193189 injected samples and it was like a sham operated control (Fig.4H). The OARSI scores in the LDN-193189-treated group were significantly lower than those in the ACLT group, even though administration of LDN-193189 was performed after the onset of disease. It should be noted, though, that less protection of cartilage was afforded, as judged by the OARSI severity scores, to the knee joints treated with LDN-193189 post-onset of OA compared to when knee joints were treated with LDN-193189 pre-onset of OA (compare Fig. 3I and Fig. 4I).

**Fig. 4.**
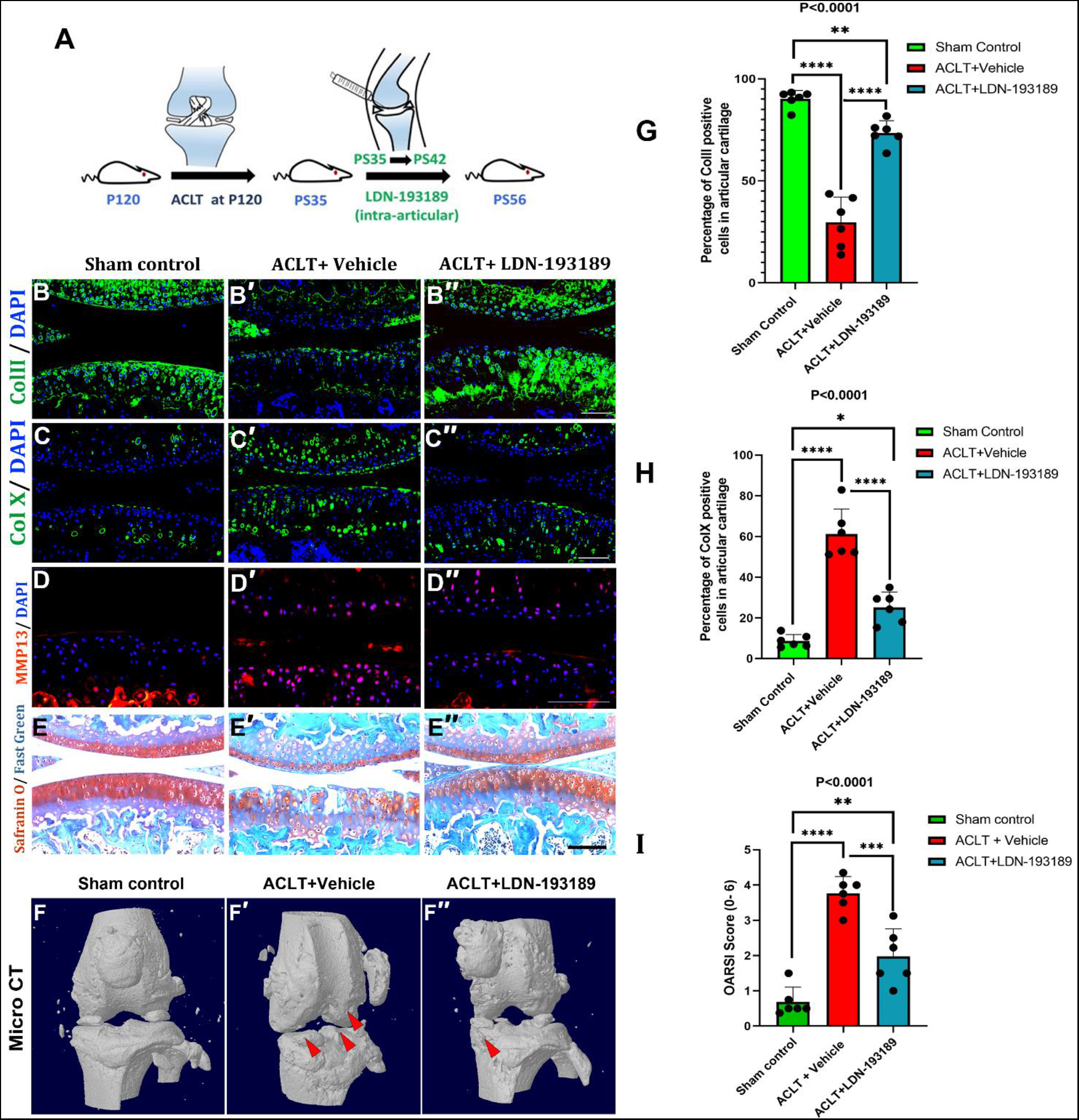
Inhibition of BMP signaling post onset of OA attenuates disease severity. **(A)** Schematic for local inhibition of BMP signaling, using of LDN-193189, in the knee joint of wildtype mouse post-surgical onset of OA. **(B-E″)** Longitudinal sections through the knee joints of sham (B-E), “ACLT + vehicle” control (B′-E′) and “ACLT + LDN-193189” (B″-E″) mice at 56 days post-surgery (PS56). Immunoreactivity for ColII (B-B″), ColX(C-C″), MMP13 (D-D″). (E-E″) Safranin O staining. (**F-F″**) 3-D rendering of μCT scan at resolution of 5.86 μm per pixel in sham, “ACLT + vehicle” control and “ACLT+ LDN-193189” injected knee joint at PS56 (Red arrows mark osteophytes). **(G)** Quantification data for ColII, one way ANOVA was performed along the three sets and p<0.0001(****). The comparison of Sham control vs ACLT+vehicle; p<0.0001 (****), Sham control vs ACLT+ LDN-193189; p=0.0088 (**) and ACLT+vehicle vs. ACLT+ LDN-193189 p<0.0001(****). **(H)** Quantification data for ColX, one way ANOVA was performed along the three sets and p<0.0001(****). The comparison of Sham control vs ACLT+vehicle; p<0.0001 (****), Sham control vs ACLT+ LDN-193189; p=0.0111 (*) and ACLT+vehicle vs. ACLT+ LDN-193189 p<0.0001(****). **(I)** OARSI score, one-way ANOVA was performed, p<0.0001(****), the comparison of means of Sham control vs ACLT+vehicle; p<0.0001 (****), Sham control vs ACLT+LDN-193189; p=0.0042 (**) and ACLT+vehicle vs. ACLT+LDN-193189 p=0.0002(***). Scale bar = 100μm, n=6 per group.

We have observed that intra-articular administration of LDN-193189 provides protection against OA-like changes at least for 14 days post injection (Fig.4). Next, we wanted to investigate the potential for clinical translatability of LDN-193189 or similar molecules as disease modifying agents. We examined whether LDN-193189 can confer longer-term protection against surgically induced OA by emulating a clinic-like regimen of minimum dosage and maximum efficacy over extended durations of time. Our data (Fig. S2) as well as the existing literature (*36*) suggest that molecular changes associated with OA are apparent within 28 days of ACLT. Hence, we conducted ACLT at P120, injected LDN-193189 intra-articularly on PS28, PS30, and PS32, and harvested the knee joint 56 days later at PS84. ColII expression (compare Fig. 5B with Fig. 5B‴) and cartilage specific proteoglycan content (compare Fig. 5D with Fig. 5D‴) were largely preserved in the LDN-193189 injected specimen when compared to the vehicle control. In addition, ColX immunoreactivity was significantly lower in LDN-193189-treated knee joints compared to vehicle-injected knee joints (compare Figs. 5C and 5C′). This set of data suggests that even after the onset of surgically induced OA, blocking the BMP signaling pathway locally can offer protection for at least 56 days in mice.

**Fig. 5.**
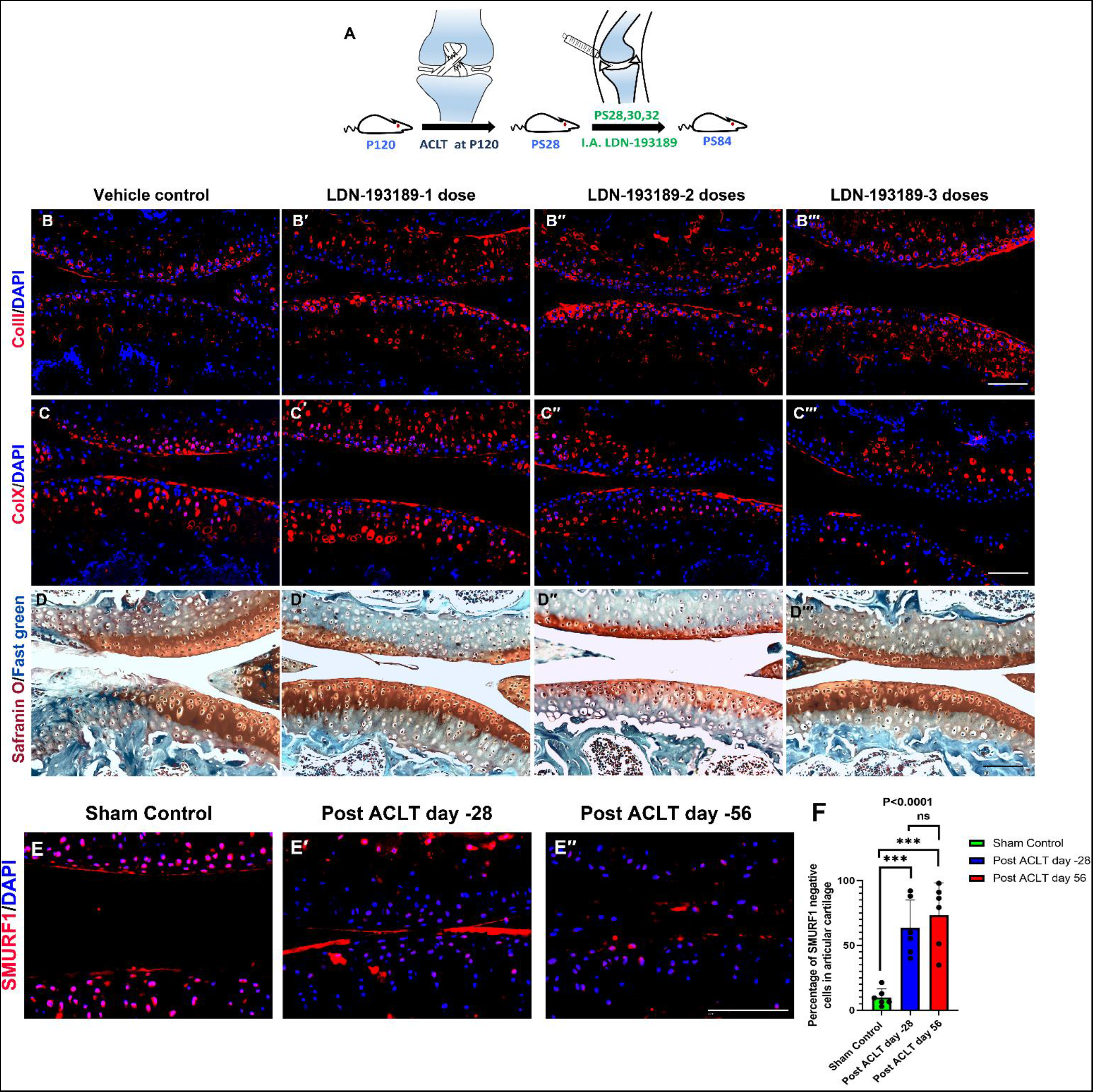
Local inhibition of BMP signaling post-onset of surgically induced OA attenuates the severity of OA associated changes for longer duration. **(A)** Schematic for local inhibition of BMP signaling, using of LDN-193189, in the knee joint of wildtype mouse post-surgical onset of OA. Longitudinal sections through the knee joints of ACLT induced OA mice (**B-D‴**). “ACLT + vehicle” control (B, C and D), “ACLT + LDN-193189 one dose” (B′, C′ and D′), “ACLT + LDN-193189 two doses” (B″, C″ and D″), “ACLT + LDN-193189 three doses” (B‴, C‴ and D‴) mice at 84 days post ACLT. Immunoreactivity for ColII (B- B‴), ColX (C- C‴) and Safranin O staining (D- D‴). **(E-E″)** Immunoreactivity for SMURF1 in Sham control (E), post ACLT 28 days (E′) and 56 days (E″). **(F)** Quantification of SMURF1 negative cells in articular cartilage, one way ANOVA was performed along the three sets with p<0.0001. The comparison of Sham control vs. post ACLT Day28 p=0.0007 (***), Sham control vs. Post ACLT Day 56 p=0.0001(***) and Post ACLT Day28 vs. post ACLT Day 56 p=0.6523 (ns). Scale bar = 100μm, n=6 per group.

### 5. Mechanistic insight into the pathogenesis of OA from a developmental biology perspective

Recently, Singh *et al.*, demonstrated that immobilisation of chick or mouse embryos results in transient cartilage differentiation at the expense of articular cartilage differentiation, which is associated with ectopic activation of BMP signaling (*23*). Further, this study also demonstrated that this ectopic activation is associated with a concurrent downregulation of expression of SMURF1, an intracellular inhibitor of the BMP signaling pathway. (*23*). SMURF1 expression was found to be lower in mouse articular cartilage 28 and 56 days after ACLT (Fig. 5E-E″). SMURF1 quantified data shows a significant decrease in SMURF1 expression at post-ACLT Days 28 and 56 (Fig. 5F) when compared to the control group. This suggests that the molecular mechanism of articular cartilage maintenance via mechanical regulation is conserved between embryonic and postnatal stages and is likely involved in pathologies such as OA.

### 6. Effect of local inhibition of BMP signaling on inflammatory responses in a surgically induced osteoarthritic mouse model

We performed an analysis for candidate inflammatory response molecules, which are known to be involved in the development of osteoarthritis(*37*, *38*). The NFkB immunoreactivity in the articular cartilage of the vehicle-treated ACLT group was significantly increased (Fig. 6B′), but it was minimal in the LDN-193189-treated or sham-operated groups (Figs. 6B″ and 6B, respectively). We also looked at TNF-immunoreactivity in osteoarthritic cartilage after LDN-19189 treatment and found that it was significantly higher in the ACLT group injected with only vehicle (Fig. 6C′, white arrow), while the LDN-19189 treated group showed minimal immunoreactivity (Fig. 6C″), and it was similar in sham-operated mice where TNF-could be detected in subchondral bone (Fig. 6C). Quantitative analysis indicates TNF-and NFkB were significantly lower in LDN-193189-treated samples compared to vehicle controls (Fig.6D & 6E, respectively). Therefore, inhibition of BMP signaling not only inhibits OA markers in articular cartilage but also reduces inflammation associated with osteoarthritis.

**Fig. 6.**
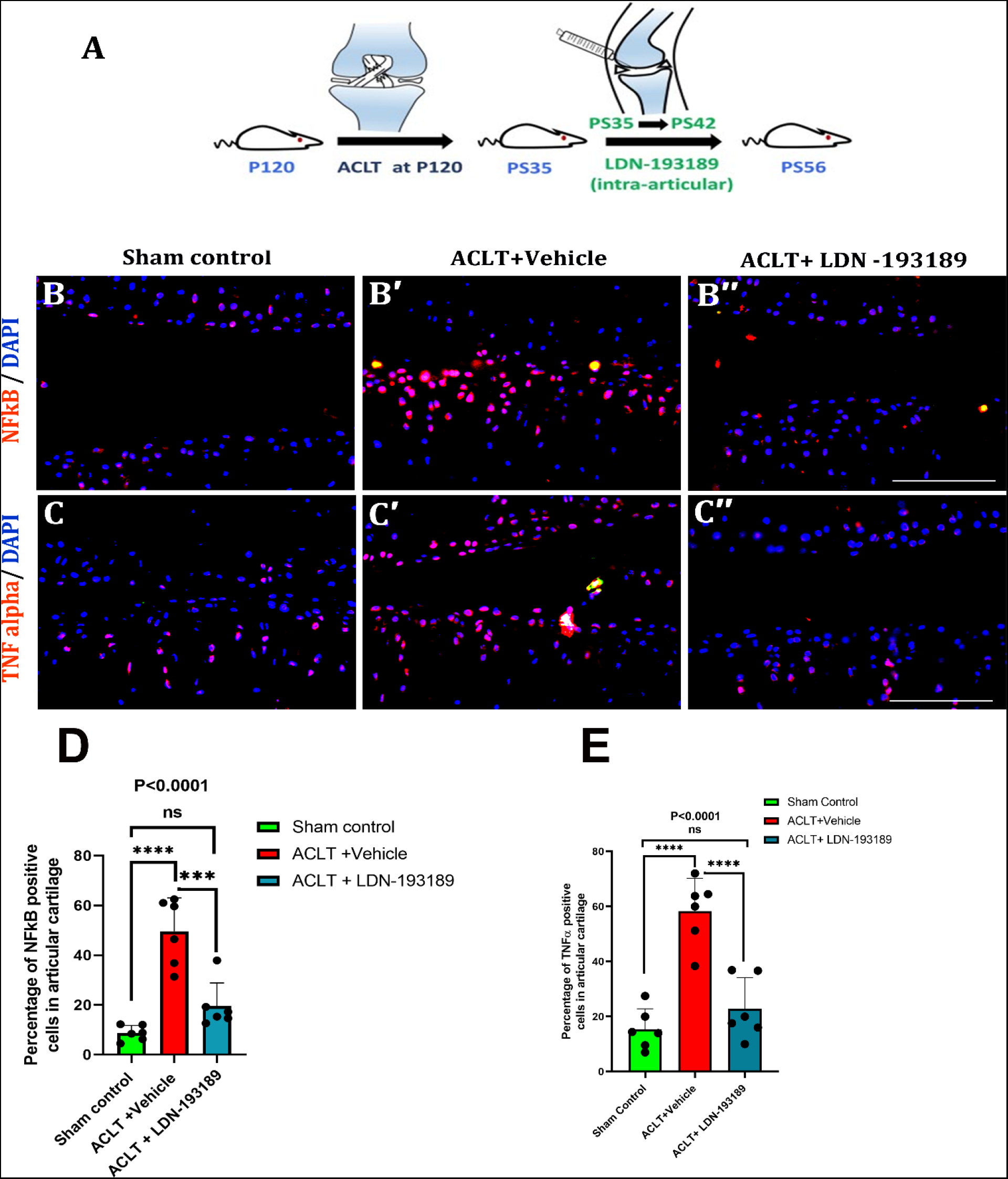
Effect of local inhibition of BMP signaling on inflammatory responses in a surgically induced osteoarthritic mouse model. **(A)** Schematic for local inhibition of BMP signaling, using of LDN-193189, in the knee joint of wildtype mouse post-surgical onset of OA. **(B-C″)** Longitudinal sections through the knee joints of sham (B-C), “ACLT + vehicle” control (B′-C′) and “ACLT + LDN-193189” (B″-C″) mice at 56 days post-surgery (PS56). Immunoreactivity for NF-κB (B-B″) and TNF-α (C-C″) levels. White arrows indicate TNF-α positive cells. **(D)** Quantification data for NF-κB, one way ANOVA was performed along the three sets and p<0.0001(****). The comparison of Sham control vs ACLT+vehicle; p<0.0001 (****), Sham control vs ACLT+ LDN-193189; p=0.1601 (ns) and ACLT+vehicle vs. ACLT+ LDN-193189 p=0.0002(***). **(H)** Quantification data for TNF- α, one way ANOVA was performed along the three sets and p<0.0001(****). The comparison of Sham control vs ACLT+vehicle; p<0.0001 (****), Sham control vs ACLT+ LDN-193189; p=0.4479 (ns) and ACLT+vehicle vs. ACLT+ LDN-193189 p<0.0001(****). Scale bar = 100μm, n=6 per group.

## Discussion

This study suggests existence of striking resemblance between the molecular changes associated with pathogenesis of osteoarthritis and endochondral ossification. Moreover, the temporal order of the expression of molecular markers appears in ACLT induced OA was found to be similar with transient cartilage differentiation i.e., endochondral bone formation. Inhibition of transient cartilage differentiation via blocking IHH signaling has been reported to inhibit and attenuate the severity of osteoarthritic phenotype post ACLT (*39*–*41*). However, so far, no IHH signaling inhibitor has been approved for clinical use. This could be because Ihh loss-of-function, during embryonic development, results only in a temporary delay in hypertrophic differentiation – an essential step of endochondral ossification, which eventually gets restored and even accelerated at postnatal stages in mice (*41*, *42*). Taken together, these studies give a crucial hint that blocking transient cartilage differentiation is a viable strategy to manage osteoarthritic changes in the articular cartilage. This hypothesis is in line with what has been suggested earlier in the literature (*43*–*45*).

BMP signaling is known to play a critical role in transient cartilage differentiation which is regulated by an intracellular BMP signaling inhibitor SMURF1.Our results indicate up-regulated BMP signaling with concomitant depletion of SMURF1 is associated with the pathogenesis of OA. Thus the low level of BMP signaling maintained by SMURF1 and divergence from it becomes inimical for cartilage health Previously, intra-peritoneal administration of a BMP signaling inhibitor, LDN-193189, has been shown to reverse the phenotype associated with *Fibrodysplasia ossificans progressive* (FOP), a condition where progressive heterotopic ossification of muscle is observed upon injury, due to constitutive activation of BMP signaling (*33*). In this study, we found that activation of BMP signaling is both necessary and sufficient for the pathogenesis of OA in mice. The necessity of BMP signaling in the onset of osteoarthritis like changes in the articular cartilage has been demonstrated using both genetic and pharmacological means whereas sufficiency has been demonstrated using genetic means. Further, analysis of patient samples suggests an association between osteoarthritis and activation of BMP signaling in the articular cartilage cells. Since we have used *TgCol2a1-Cre-ERT2* mediated recombination as the means to activate expression of constitutively activated BMP receptor (caBMPRIA) we cannot rule out the possibility that BMP signaling has also been activated in the growth plate cartilage of adult mice and the molecular and cellular changes observed is partly due to activated BMP signaling in the growth plate cartilage. However, all our experiments have been done after skeletal maturity of mice so there are least contribution of observed phenotype due to change in the growth plate chondrocyte. Moreover, the changes were first observed in the superficial layers of articular cartilage suggesting that the changes observed were primarily due to ectopic activation of BMP signaling in the articular cartilage.

Interestingly, we also observed proliferation in articular cartilage cells, as assessed by enhanced BrdU uptake, post ACLT or activation of BMP signaling. Our data suggests that articular cartilage cells, originally having low regenerative potential and proliferative capacity, display a regenerative response upon ACLT or upregulation of BMP signaling. However, this leads to an altered tissue microenvironment that promotes transient cartilage differentiation at the expense of articular cartilage. Thus, instead of healing by regeneration it further promotes the disease condition. Prophylactically blocking BMP signaling *in situ* using LDN-193189 led to an attenuation in the severity of osteoarthritic phenotype following surgical induction in mice. Further, our investigation suggests that administration of LDN-193189 after the onset of OA not only halts the progression of OA but also an intense Safranin O-stained cartilage tissue appears which is negative for transient cartilage markers, suggesting that new cartilage formation takes place. A recent study published by Liu et *al.* in 2020 also suggests the role of BMP signaling inhibition to target osteoarthritis. However, the inhibitor has been given intra-peritoneally which is not a feasible option for patients due to its global consequences on the body.

Finally, it has to be acknowledged that while transient cartilage differentiation could be involved in the initiation of the disease, the inflammation associated with OA can determine the severity and course of disease progression (*46*, *47*). Despite a large body of existing literature, there is no clear demonstration of the hierarchy between the onset of inflammation and transient cartilage differentiation. Literature suggest that BMP signaling regulates endothelial inflammatory pathway after cardiac ischemic injury (*48*). Our study also signifies that pharmacologically blocking BMP signaling in surgically induced OA also prevents inflammatory response activation. However, whether BMP signaling directly regulates inflammatory pathway or it induces chondrocyte hypertrophy causing inflammation due to altered joint mechanics, further needs to be investigated. Nonetheless our study demonstrates that *in situ* inhibition of BMP signaling, and consequently transient cartilage differentiation, may be a potent means of disease-modifying therapy for osteoarthritis.

## Acknowledgements

We are immensely grateful to Prof. YiPing Chen at Tulane University, USA, for the gift of mouse strains. We thank Prof. Frank Beier of Western University, Ontario, Canada for teaching APJ the method of ACL transection. We sincerely thank Shuchi Arora and Ankita Jena for their critical comments on the manuscript. We are highly grateful to Niveda Udaykumar and Saahiba Thaleshwari for their help in blind OARSI scoring. We thank Mr. Naresh Gupta for assistance with mouse experiments.

## Author’s contribution

A.B., A.P.J. and B.K. designed the experiments and A.P.J., B.K., A.K.S. S.V.N. and S.F.I. conducted experiments, collected, and analysed data. A.P.J., B.K. and S.F.I. prepared the manuscript; N.A. conducted the cell-based LDN-193189 assay. A.B., C.D.B., A.J.R. edited the manuscript along with A.P.J.; B.K. and A.K.S. provided the data for inflammation response studies and mechanistic data including Smurf expression analysis; A.J.R. and A.H.K.R. collected and analysed human cartilage samples; H.W. and S.A. performed the scoring for osteoarthritis

## Competing Interests

The authors declare the following competing interests:

The use of BMP inhibitors as locally administered agents using sustained drug delivery vehicle(s) has been submitted for patent via Indian patent application number – **201911044840**.

The inventor(s) are:

1. Dr. Amitabha Bandyopadhyay–IIT Kanpur, India,
2. Mr. Akrit Pran Jaswal -IIT Kanpur, India.
3. Mr. Bhupendra Kumar-IIT Kanpur,India.
4. Dr. Praveen Vemula – Institute for Stem Cell Biology and Regenerative Medicine, India.
5. Dr. Manohar Mahato - Institute for Stem Cell Biology and Regenerative Medicine, India.

