## Supplementary material for "BMP signaling: A significant player and therapeutic target for osteoarthritis": no

<u>**Supplementary materials (SM)**</u>

### Detailed methodology

#### Study design:

**Sample size:** We used statistical power calculations using G*Power3.1 software to generate the sample size from previously conducted studies in mice using Bmp2/4 DCKO mice (24).

**Data exclusion:** No data was excluded from the end study. Rules for stopping data collection: Data collection was stopped at end points pertinent to different experiments since tissues were harvested at the end of each experiment. The relevant time points were determined for each experiment based on similar experiments reported previously as well as from developmental biology insights and no outliers were present or reported. Replication of data was ensured for all experiments including protocols involving classical histology/ molecular histology by repeating multiple times to ensure reproducibility of data.

**Experimental design:** The study was a controlled laboratory experiment involving animal subject, human samples, and cell culture studies. Primary nature of data collected, recorded, and analysed was imaging data, viz. microscopic imaging, micro-CT imaging data and statistical software used accordingly.

**Randomization:** Age matched, gender matched, genotype matched animals were used to minimize randomization in experiments. For all genetic induction experiments 2 groups were used: 1) Vehicle injected and 2) tamoxifen injected. In experiments involving surgical induction with or without genetic induction, 3 groups were used: 1) Sham surgical control 2) ACLT + vehicle 3) ACLT + Tamoxifen / LDN-191389 treated, depending on the type of experiment.

**Blinding:** OARSI scoring were performed by researchers as OARSI guideline, and they were not involved in the animal experiments and who were blinded to the treatment groups**.** Data was then unblinded and analysed by the researchers carrying out the experiments. Details for animal and human subjects are provided in the sections following this.

**Genotyping**

Mice were genotyped for the presence of caBmpr1a using PCR with primers 5′-CGAAGATATGCGTGAGGTTGTGT-3′ and 5′- TGGGCGAGAGGGGAAAGAC - 3′ while *TgTgCol2a1-Cre-ERT2* transgene was genotyped using primer pair 5′-CACTGCGGGCTCTACTTCAT-3′ and 5′- ACCAGCAGCACTTTTGGAAG- 3′. *ROSA26 mT/mG* was genotyped using primers 5′-CTCTGCTGCCTCCTGGCTTCT-3′ and 5′- TCAATG GGCGGGGGTCGTT - 3′. Bmp2c/c was genotyped using 5′-GTGTGGTCCACCGCATCAC-3′ and 5′- GGCAGACATTGTATCTCTAGG - 3′ and Bmp4c/c was genotyped using 5′-AGACTCTTTAGTGAGCATTTTCAAC-3′ and 5′-AGCCCAATTTCCACAACTTC - 3′ primer pairs.

**ACLT Surgery**

ACLT was performed in left hind limb of all mice. Patella was dislocated medially to expose femorotibial joint cavity and ACL was transected using a micro-surgical blade under the microscope. After ACL transection, patella was repositioned and quadriceps femoris muscle was sutured to prevent its dislocation. In the sham group, the joint cavity and ACL were exposed but left intact and sutured. Surgical incision was sealed using Vicryl 5-0 Absorbable suture. Povidone-Iodine solution was applied over the sutured skin to prevent microbial infections.

**BMP signaling gain-of-function**

*pMes-caBmpr1a;TgTgCol2a1-Cre-ERT2* male mice were injected with three consecutive doses of tamoxifen (Sigma-Aldrich, 2.5 mg/25 g body weight) starting from P70 to induce Cre mediated recombination and harvested at post TAM injection day 7,14, 21, 28, 56 and 6 months for molecular and histological analysis.

**BMP signaling loss-of-function**

All animals used for BMP signaling loss-of-function were homozygous males for the conditional alleles of *Bmp2* and *Bmp4*. Three doses of tamoxifen (Sigma-Aldrich, 2.5 mg/25 g body weight) were injected in *Bmp2*$^{c/c}$*; Bmp4*$^{c/c}$; *TgTgCol2a1-Cre-ERT2* mice intraperitoneally starting from P70 for three consecutive days to activate Cre-mediated recombination to deplete BMP2 and BMP4 in adult cartilage. pSMAD1/5/9 immunoreactivity was used to assess the status of BMP signaling before/after ACLT and/or with/without TAM-induced recombination.

Local inhibition of BMP signaling was achieved through administration of 6μl solution of 10μM LDN-193189, prepared in 3% 2-(hydroxypropyl)-β-cyclodextrin (w/v in PBS) in the knee joint cavity. Local inhibition of BMP signaling was done at two different time points. In one set, LDN-193189 was injected daily from 14$^{th}$ to 21$^{st}$ day post ACLT. In the other set LDN-193189 was administered daily from 35$^{th}$ to 42$^{nd}$ day post ACLT.

To observe the persistent effect of LDN-193189 over a prolonged period as well as to ascertain the optimum number of doses needed for providing long term protection following ACLT, LDN-193189 was injected on post-surgery day 28 (single dose) or post-surgery days 28 and 30 (two doses) or post-surgery day 28, 30 and 32 (three doses). Tissues were harvested at post ACLT day 84 for further analysis. In all experiments the vehicle control animals were injected with 3% 2-(hydroxypropyl)-β-cyclodextrin (w/v in PBS).

**Determination of the recombination frequency at P70 in mice articular cartilage.**

For determination of recombination frequency under the *Col2a1Cre, TgCol2a1::Cre-ERT2;ROSA26::TdTomato-EGFP* compound mice line has been used. Tamoxifen was injected at P70 along with vehicle control group. At post TAM injection day 14, animals were harvested, tissue sectioned and anti GFP antibody was performed. Total number of GFP positive cells in articular cartilage were calculated in test v control sample. 75.50 % percent of articular cartilage cells were positive for anti GFP antibody which results 75.50% recombination frequency of articular chondrocyte at P70 (Fig. S4).

**Cell proliferation analysis**

Cell proliferation status was assessed using BrdU incorporation assay following intraperitoneal delivery of 100mg BrdU/10ml PBS/kg of body weight for three consecutive days prior to the day of harvesting tissues.

**LDN-193189 activity assay**

BRITER (BMP Responsive Immortalized Reporter) cell line(*25*) were used to determine optimum concentration of LDN-193189 (Sigma Aldrich, cat.no SML0559) required to inhibit BMP signaling.

**Tissue Processing, histology, and immunohistochemistry**

Post harvesting, tissues were fixed in 4% paraformaldehyde for 16h at 4°C and then washed in PBS for 2h. The tissues were decalcified in 14% EDTA solution (pH 7.4) for 28 days with regular changes of EDTA solution at least once in a week. Decalcified tissues were embedded in paraffin wax and sectioned at 5 µm thickness. For immunohistochemistry, sections were treated with xylene and then rehydrated using descending alcohol gradient. Antigen retrieval was done using citrate buffer except for collagens. ColII and ColX antigens were retrieved by treatment of acidic solution of 0.05% pepsin at 37ºC for 25 minutes. Tissue sections were washed with PBS and treated with 3% $H_2O_2$ for 15 minutes to block endogenous peroxidase activity. After blocking in 5% HINGS (Heat-inactivated Goat Serum) solution for an hour, tissue sections were incubated with primary antibodies against Type X Collagen (DSHB, Cat.no. X-AC9-s, 1:20), Type II Collagen (DSHB, Cat.no. CIIC1-s, 1:20), SMURF1 (Santa Cruz Biotechnology, Cat.no.SC100616,1:100), NF-κB (CST, Cat.no.D14E12,1:400, MMP-13 (Santa Cruz Biotechnology, Cat.no.SC30073,1:100), pSMAD1/5/9 (CST Cat.no.13820S, 1:100), GFP (Invitrogen, Cat no. A6455, 1:100), BrdU (Sigma, Cat.no. B8434) and TNF-α (Santa Cruz Biotechnology, Cat.no. SC-52746, 1:100) for 16 hours at 4ºC. Tissue sections were washed with PBST and incubated with fluorescence labelled secondary antibodies for overnight at 4ºC. Secondary antibody against BrdU was conjugated with HRP (horseradish peroxidase) and detected by DAB (3,3 Diaminobenzidine).

Histological analysis was performed using Safranin O staining following standard protocol. For Safranin O/ Fast green staining, rehydrated tissue sections were treated with haematoxylin solution for 10 minutes, washed in running tap water, and treated with 0.5% Fast Green solution for 2 minutes. Thereafter, sections were dipped in 1% acetic acid and finally stained with 0.5% Safranin O solution for 2 minutes. Stained sections were dehydrated with 95% and 100% alcohol respectively, cleared with Xylene and mounted in DPX. All images were taken using Leica DM 5000B compound microscope and processed using manufacturer provided software suite(s). Human cartilage sections were stained by immunohistochemistry for pSmad1/5/9 using standard protocols (48). Briefly, antigen retrieval was performed by incubation for 1 hour at 65ºC in antigen unmasking citrate buffer solution (pH 6, Vector Laboratories, UK), after which sections were incubated with phosphatase (CST Lambda Phosphatase kit, New England Biolabs, #P0753) according to manufacturer's instructions, or incubated with buffer only. Sections were then incubated with an antibody against pSmad1/5/9 (Millipore, cat no. ab3848, at 1:50 (10 µg/ml)) followed by biotinylated goat anti-rabbit IgG (Vector Labs, cat no. BA-1000, at 1:200). Stained sections were imaged using Zeiss Axioskop 40 (Zeiss) with Progress XT Core 5 colour digital camera and ProgRes CapturePro 2.9.0.1 software (JenOptik, Germany).

**OARSI scoring for histological sections**

In our experiments, five sections were taken from different depths of the cartilage tissue and evaluated after safranin O staining. Based on Safranin O staining, changes in articular cartilage were scored blindly from 0–6 by at least two individuals who were not involved in the experiment. Every data point illustrates one animal, which is an average score from two individuals. A score of 0-2 indicates only minor changes, whereas a score of 3-6 represents significant cartilage erosion.

**Generation of quantitative data**

Quantification of ColX, ColII, NFkB and TNF alpha was done by counting percentage of articular chondrocyte positive for respective molecules. The quantification of SMURF1 was done by calculating percentage of negative cells for SMURF1 in articular cartilage.

**3-D volumetric projections of µCT data**

3-D volumetric projections were generated using CTVol 2.0 software. The scanner was set to operate at a voltage of 50 kVp, a current of 200 µA and the resolution of the images was 5.86 µm per pixel. µCT for BMP signaling gain-of-function specimens was performed, processed, and analysed at the Institutional facility of Western University, Ontario, Canada using eXplore Locus GE scanner. The resolution of the image is 40 µm per pixel and 3-D isosurface projection was generated using Parallax Innovations MicroView 2.5.0- rc15 (2.5.0-3557)

### Supplementary figure legends

**Fig. S1. Molecular characterization of mouse knee joint post ACLT**

**(A-D‴)** Longitudinal sections through the knee joints of sham (A, C), 7 days (B, D), 14 days (B′, D′), 21 days (B″, D″), 28 days (B‴, D‴) post ACLT. **(A-B‴)** Immunoreactivity for pSMAD1/5/9. **(C-D‴)** BrdU uptake. Insets display the higher magnification view of the marked regions. Scale bar = 100µm, n=5 per group

**Fig. S2. BMP signaling induced transient cartilage differentiation is necessary for the pathogenesis of OA.**

**(A)** Schematic representation depicting the generation of *$Bmp2^{c/c}$; $Bmp4^{c/c}$; TgCol2a1-Cre-ERT2* and the regimen for depletion of BMP signaling by administration of tamoxifen followed by ACLT. **(B-F″)** Longitudinal sections through the knee joints of sham (B-F), “ACLT + vehicle control” (B′-F′) and “BMP depletion + ACLT” (B″-F″) mice at 28 days post-surgery (PS28). Immunoreactivity for pSMAD1/5/9 (B-B″), ColII (C-C″), ColX (D-D″), MMP-13 (E-E″). **(F-F″)** Safranin O staining. n=5 per group. Scale bar = 100µm.

The panels where *Bmp2/4* depleted animals were subjected to ACLT are marked as “BMP depletion + ACLT”. Vehicle injected animals were used as genotype controls (“ACLT + Vehicle”. Scale bar = 100µm, n=5 per group

**Fig. S3. Determination of optimum LDN-193189 concentration to inhibit BMP signaling using BRITER (BMP Responsive Immortalized Reporter) cell line.**

Normalized relative luciferase activity after 3h of LDN-193189 treatment with different concentration (100nM, 200nM, 400nM) to inhibit BMP signaling in BRITER cell line (Fig.S3). p-value and statistical significance: The two-tailed p-value is less than 0.0001.

**Fig. S4. Determination of the recombination frequency at P70 in mice articular cartilage.**

The *TgCol2a1::Cre-ERT2;ROSA26::TdTomato-EGFP* compound mouse line has been used to study the frequency of recombination under the *Col2a1Cre* system. Tamoxifen and equivalent vehicle groups were administered at P70, and tissues were harvested at post Vehicle/ TAM injected day 14. (A) Post vehicle injected day 14, (A′) Post TAM injected Day 14. Scale bar = 100μm, n=4 per group

### Supplementary figure

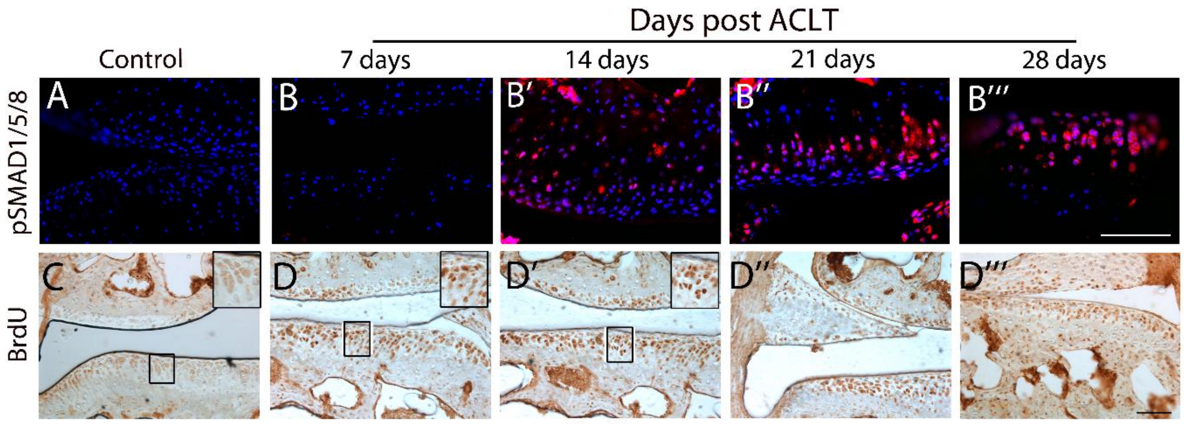


Fig. S1

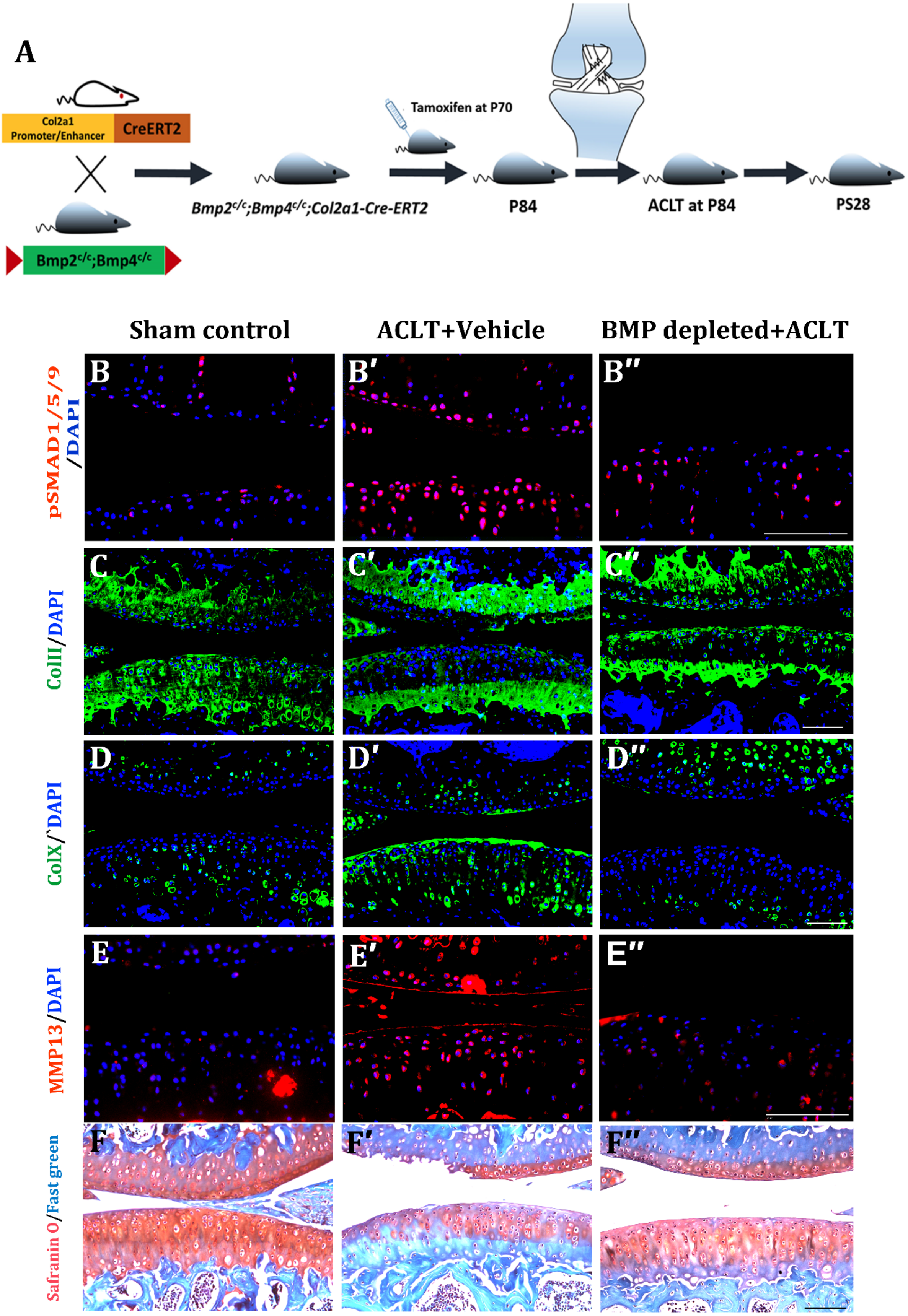
A
Col2a1 Promoter/Enhancer
CreERT2
Bmp2c/c;Bmp4c/c
Tamoxifen at P70
Bmp2c/c;Bmp4c/c;Col2a1-Cre-ERT2
P84
ACLT at P84
PS28
Sham control
ACLT+Vehicle
BMP depleted+ACLT
pSMAD1/5/9/DAPI
ColII/DAPI
ColX/`DAPI
MMP13/DAPI
Safranin O/Fast green
B
B′
B″
C
C′
C″
D
D′
D″
E
E′
E″
F
F′
F″

Fig.S2

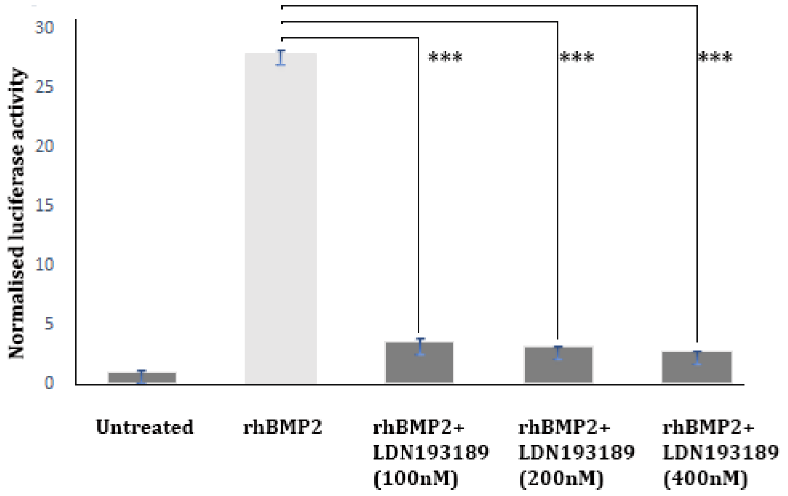

Normalised luciferase activity
0
5
10
15
20
25
30
***
***
***
Untreated
rhBMP2
rhBMP2+ LDN193189 (100nM)
rhBMP2+ LDN193189 (200nM)
rhBMP2+ LDN193189 (400nM)


Fig.S3

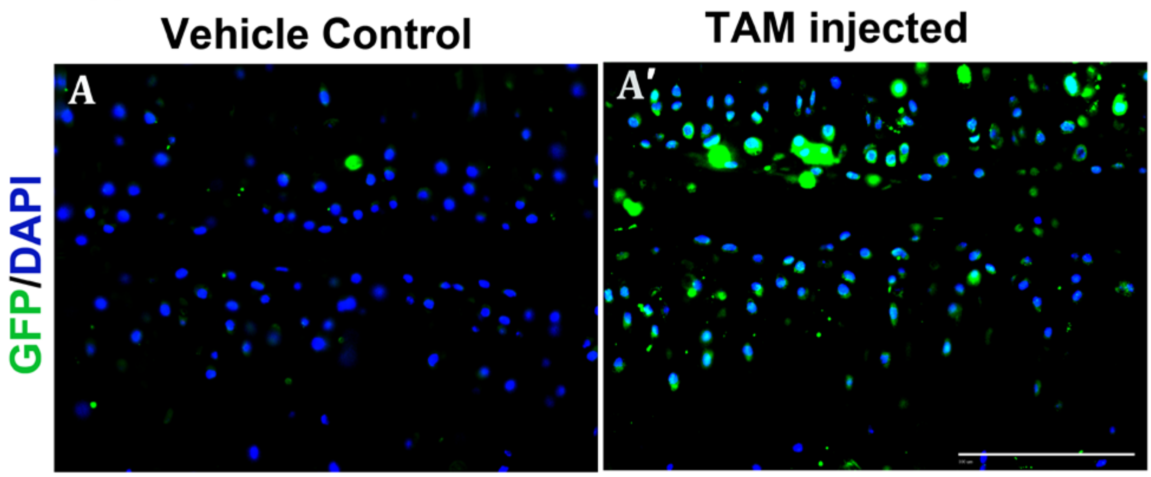

Vehicle Control
TAM injected
A
A′
GFP/DAPI


Fig.S4
